# Dark nanodiscs as a model membrane for evaluating membrane protein thermostability by differential scanning fluorimetry

**DOI:** 10.1101/2023.05.08.539917

**Authors:** Jazlyn A. Selvasingh, Eli Fritz McDonald, Jacob R. Mckinney, Jens Meiler, Kaitlyn V. Ledwitch

## Abstract

Measuring protein thermostability provides valuable information on the biophysical rules that govern structure-energy relationships of proteins. However, such measurements remain a challenge for membrane proteins. Here, we introduce a new experimental system to evaluate membrane protein thermostability. This system leverages a recently-developed non-fluorescent membrane scaffold protein (MSP) to reconstitute proteins into nanodiscs and is coupled with a nano-format of differential scanning fluorimetry (nanoDSF). This approach offers a label-free and direct measurement of the intrinsic tryptophan fluorescence of the membrane protein as it unfolds in solution without signal interference from the “dark” nanodisc. In this work, we demonstrate the application of this method using the disulfide bond formation protein B (DsbB) as a test membrane protein. NanoDSF measurements of DsbB reconstituted in dark nanodiscs show a complex biphasic thermal unfolding pattern in the presence of lipids with a minor unfolding transition followed by a major transition. The inflection points of the thermal denaturation curve reveal two distinct unfolding midpoint melting temperatures (T_m_) of 70.5 °C and 77.5 °C, consistent with a three-state unfolding model. Further, we show that the catalytically conserved disulfide bond between residues C41 and C130 drives the intermediate state of the unfolding pathway for DsbB in a nanodisc. We introduce this method as a new tool that can be used to understand how compositionally, and biophysically complex lipid environments drive membrane protein stability.

## Introduction

Membrane proteins represent a class of proteins that are challenging targets for biophysical studies because they associate with unique asymmetric lipid environments. Evaluating the thermal stability of membrane proteins relies on spectroscopic or calorimetric assays that perform well and are robust for water-soluble proteins such as circular dichroism (CD)^1^, differential scanning calorimetry (DSC)^2^, and differential scanning fluorimetry (DSF)^3^. These tools often fall short for membrane proteins because they require a model membrane system for *in vitro* studies, which often convolute the signal and measurement. Detergent micelles are the most widely used model membrane systems for biophysical studies, despite their known effect on membrane protein structure, stability, and function^4,5^. To date, few studies tabulated in the membrane protein thermodynamic database (MPTherm) report thermostability measurements for membrane proteins in the presence of lipids^6^ despite the key role lipids play in thermal stability^7-9^. This is largely due to technical hurdles associated with developing simple tools to investigate the effects of lipids on membrane protein stability.

Model membrane systems used for membrane protein stability measurements range from micelles and bicelles to more native-like systems such as nanodiscs^10^, saposin-derived lipid nanoparticles^11^, styrene-maleic acid copolymers^12^, and proteoliposomes^13^. Many sophisticated attempts have been made to evaluate how lipids impact membrane protein stability using a variety of approaches. For example, an ion mobility mass spectrometry approach showed that specific lipid types increased protein unfolding resistance for several membrane protein systems when compared to micelle conditions alone^14^. Nji *et al*. developed a thermal shift assay to probe the effect of lipid interactions on membrane protein stability^15^. That work revealed that cardiolipin increased the thermal stability of a sodium–proton antiporter when added to the detergent-solubilized membrane protein sample. Treuheit *et al*. was the first to study the thermal stability of a membrane-associated protein, cytochrome P4503A4 (CYP3A4), in a nanodisc by DSC^16^. The results showed an increase in the thermal stabilization of CYP3A4 incorporated into nanodiscs compared to an aggregated state in solution. Flayhan *et al*. tested the use of saposin lipid nanoparticles (SapNPs) as a model membrane system for membrane protein reconstitutions^17^. Three different membrane protein systems were reconstituted in SapNPs, two integral membrane peptide transporters (DtpA and PepT) and the small-conductance mechanosensitive channel T2, for thermostability measurements by nano-format of differential scanning fluorimetry (nanoDSF). All three membrane protein test systems revealed an increase in thermal stabilization (i.e., melting temperature) in SapNPs compared to n-dodecyl-β-D-maltoside (DDM) micelles. These attempts suggest we still lack robust tools for evaluating and quantitatively comparing how lipids regulate the structure and stability of membrane proteins.

Recently, a membrane scaffold protein (MSP) construct was engineered that replaces all tryptophan and tyrosine residues with phenylalanine for reconstituting membrane proteins in nanodiscs making them fluorescently dark^18^. Nanodiscs are attractive systems because the lipid composition can be precisely controlled and therefore, can be used as a model membrane to systematically dissect the effect of lipid types on membrane protein thermal stability. In this work, we introduce a new method that couples the non-fluorescent MSP to form dark nanodiscs with nanoDSF measurements of intrinsic tryptophan fluorescence. This offers a facile measurement of the thermal unfolding of reconstituted membrane proteins without signal interference from the nanodisc. Further, thermostability measurements by nanoDSF offer many advantages in that it is a label-free approach that monitors the shift in intrinsic tryptophan fluorescence as the protein unfolds and does not require a high sample consumption. We test the utility of this approach using the *E. coli* disulfide bond formation protein B (DsbB) as a model membrane protein system. The topology of DsbB is shown in **Figure 1** and consists of four transmembrane domains (TMD1–TMD4) with a catalytically-conserved interloop disulfide bond that plays an important part in the reaction cycle^19^. We use this as a model membrane protein because the thermodynamic behavior of DsbB has been extensively evaluated under chemical denaturation conditions making it an ideal test system for comparison^20,21^.

**Figure 1.**
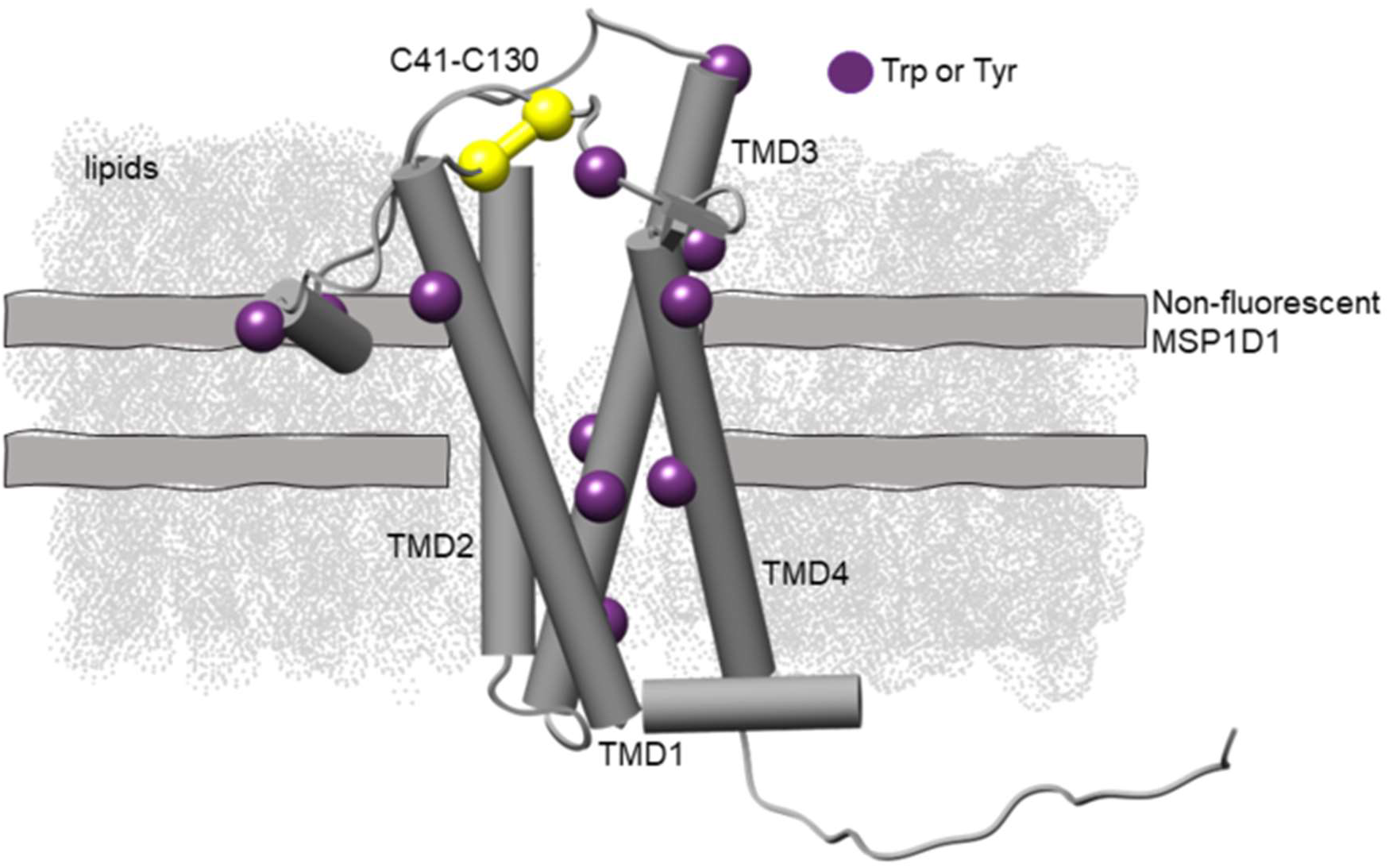
The structural topology and location of the tryptophan and tyrosine residues mapped onto the NMR-derived model of DsbB reconstituted in a dark nanodisc (PDB ID 2K73). DsbB is reconstituted in a dark nanodisc system loaded with DMPC and DMPG lipids at a 3:2 molar ratio and the non-fluorescent MSP. The four transmembrane domains are labeled as TMD1-TMD4 and the interloop disulfide bond C41-C130 is shown in yellow. The C41-C130 interloop disulfide bond connects the two periplasmic loops. We also show the tryptophan and tyrosine residues as purple spheres for visualization.

Early studies by Otzen *et al*. combined fluorescence-based stopped-flow kinetic experiments and chemical denaturation in sodium dodecyl sulfate and dodecyl maltoside (SDS-DM) mixed micelles to study the unfolding/refolding of DsbB. These experiments revealed a three-state unfolding model for DsbB consisting of the native state in DM, an unfolding intermediate, and the SDS-denatured state^20^. The authors also performed the same experiments under reducing conditions to investigate the role disulfide bonds play in the stability of DsbB. They showed that under reducing conditions unfolding rate constants increased and that the disulfide bonds contribute to the stability of the protein. Regardless, it remains unclear if the observed intermediate state in SDS-DM mixed micelles is biophysically-relevant. In this work, we report nanoDSF thermostability data for DsbB reconstituted in our dark nanodiscs to showcase the utility of our method. Our data suggests DsbB undergoes a three-state/biphasic unfolding process in dark nanodiscs. We compare these results to a panel of different detergent micelle conditions and under conditions where the C41-C130 interloop disulfide bond is disrupted by reducing conditions.

## Results

### The thermal unfolding curve for DsbB reconstituted in a dark nanodisc follows a biphasic unfolding pattern with a minor transition followed by a second major transition

One of the most widely used proxies for overall protein stability is thermostability. We reconstituted DsbB in dark nanodiscs loaded with a 3:2 molar ratio of 1,2-dimyristoyl-sn-glycero-3-phosphocholine (DMPC) and 1,2-dimyristoyl-sn-glycero-3-phosphorylglycerol (DMPG) lipids and measured the thermostability of the protein by nanoDSF. **Figure 2A** shows the intrinsic tryptophan fluorescence ratio at 350 and 330 nm for DsbB (F350/F330 ratio) over a thermal ramp from 25 °C to 95 °C. The inflection points of the unfolding curve determined from the experimental derivative correspond to the unfolding midpoint melting temperatures (T_m_). **Figure 2B** shows the first derivative of the F350/F330 ratio as a function of temperature. **Figure 2C** shows a representative SDS-PAGE gel demonstrating sample purity following a reconstitution reaction for DsbB in a dark nanodisc. The experimental derivative of the unfolding curve reveals two distinct T_m_ values of 70.5 °C and 77.5 °C. These results suggest that DsbB in a nanodisc follows a three-state/biphasic unfolding model: native (**N**) **→** intermediate (**I**) **→** unfolded (**U**). To confirm that the nanodisc does not contribute to the three-state/biphasic unfolding mechanism, we performed nanoDSF measurements on empty dark nanodiscs (i.e., in the absence of protein) loaded with the same lipid content. Indeed, the dark nanodisc itself is fluorescently undetectable over the temperature gradient (**Figure 2A, grey line**). This can be compared to the results for empty nanodiscs prepared using conventional (fluorescent) MSPs (**Figure 3A-B**). The near-baseline signal for the empty dark nanodisc sample up to roughly 85 °C suggests that the engineered MSP used for reconstitutions does not interfere with the signal observed for the target DsbB protein. Furthermore, DMPC and DMPG lipids have melting temperatures below the range of the thermal gradient measured (i.e., below 25 °C) and both are cylindrical lipids that lack the required geometry for a transition to a hexagonal or inverted hexagonal phase^22^. Therefore, we do not expect that the phase transition of the lipids used in this work contributes to the unfolding transitions observed for DsbB. We also report the fraction unfolded curves (baseline corrected) for visualization of the T_m_ values at the inflection point which is shown in **Figure S1A-D**. We also report the dynamic light scattering measurements collected in **Figure S2A-E (third row)** as a cumulative radius plot to evaluate any contributions of aggregation to our unfolding curve data.

**Figure 2.**
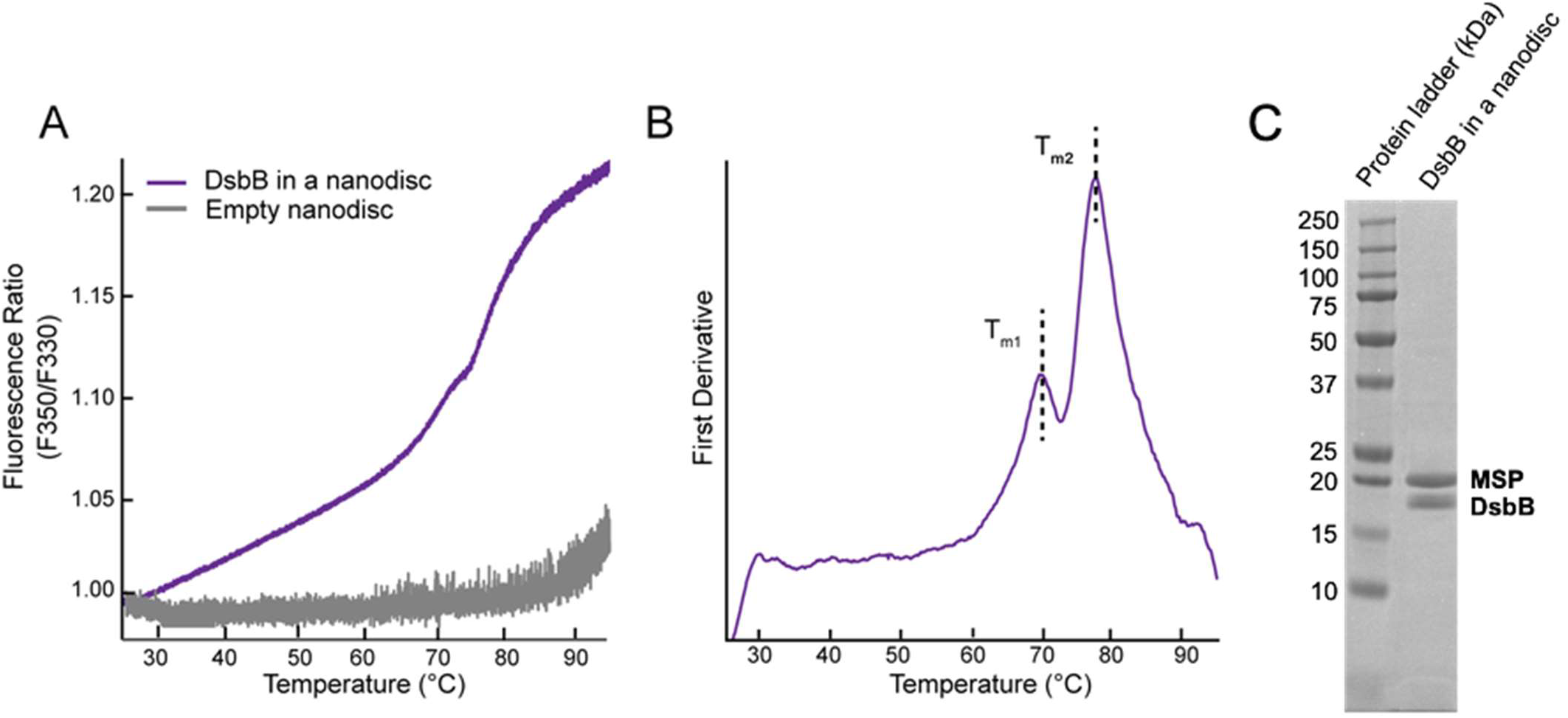
NanoDSF thermal unfolding curve for DsbB in a dark nanodisc. (**A**) The F350/F330 thermal unfolding curve for DsbB reconstituted in a dark nanodisc model membrane system (violet). The thermal unfolding curve for empty dark nanodiscs is shown in grey. (**B**) The first derivative plot of the F350/F330 ratio with respect to temperature for DsbB in a nanodisc. The inflection points correspond to T_m_ values of 70.5 °C and 77.5 °C, respectively. All samples were run in triplicate for three independent preparations and the average is plotted. (**C**) Representative SDS-PAGE gel following size exclusion to showcase the successful reconstitution of DsbB in a dark nanodisc.

**Figure 3.**
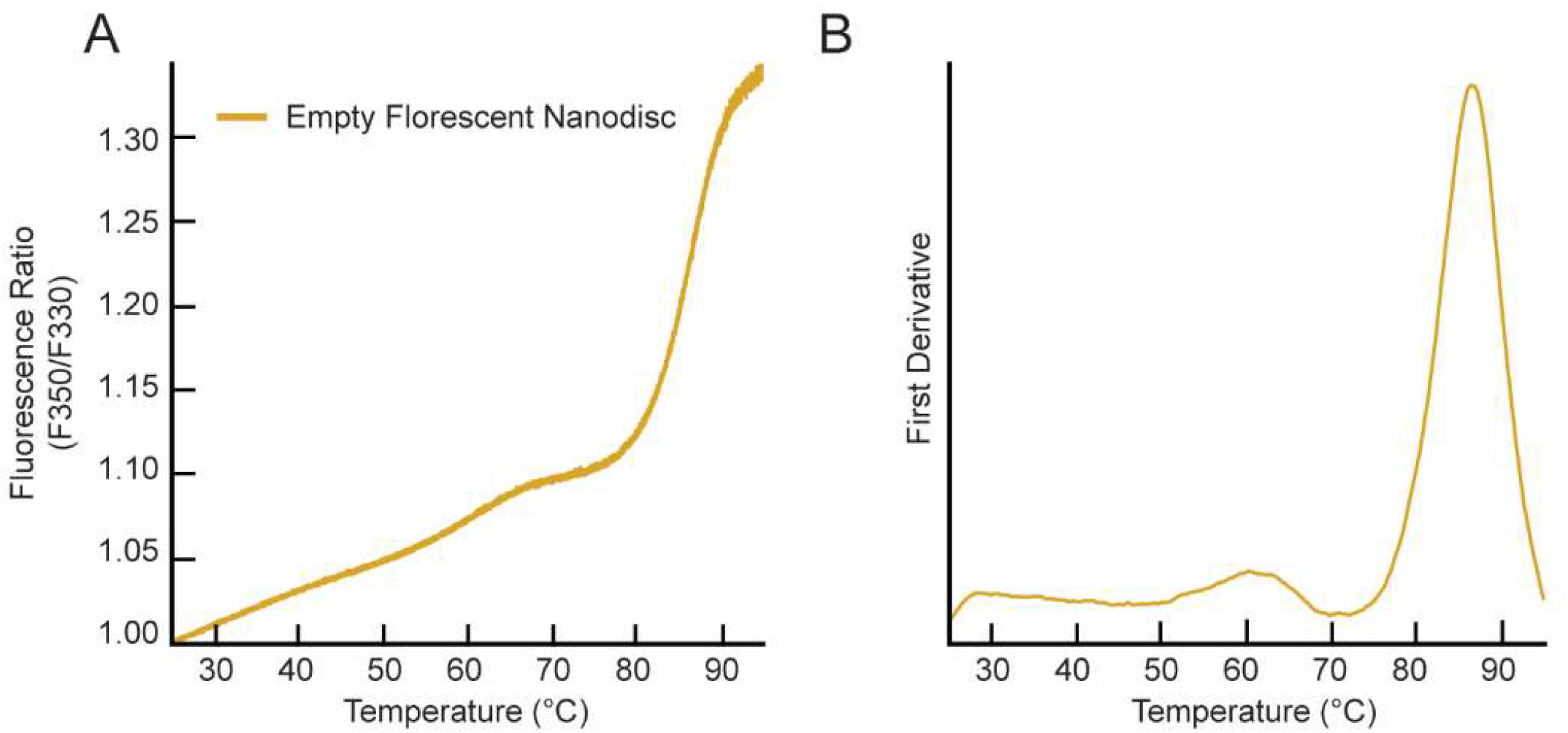
NanoDSF measurement for the empty fluorescent MSP-based nanodisc. (**A**) The F350/F330 thermal unfolding curve for the empty fluorescent MSP-based nanodisc. (**B**) The first derivative plot of the unfolding curve shows an inflection point at a T_m_ value of 86.6 ± 0.6 °C. All samples were run in triplicate and the average is plotted.

### The thermal unfolding curves for DsbB under a panel of detergent micelle conditions each show a major unfolding transition and decreased T_m_ values compared to nanodiscs

We selected a range of detergents with different physiochemical properties and hydrocarbon chain lengths to compare the thermostability of DsbB in nanodiscs to different micelle conditions. **Figure S3A-E (first row)** shows the F350/F330 ratio for the unfolding curves for detergent-solubilized DsbB in DDM, lauryl maltose neopentyl glycol (LMNG), n-decyl-β-maltoside (DM), lauryldimethylamine oxide (LDAO), and lyso-myristoylphosphatidylglycerol (LMPG) micelles. **Figure S3A-E (second row)** shows the respective experimental derivative plots for the unfolding curves under each detergent micelle condition revealing a single major inflection point corresponding to a single T_m_ value. Detergent-solubilized DsbB switches to a single major unfolding transition in the case of DDM and LMPG micelles. A second less-defined peak can be seen towards the end of the thermal gradient, roughly 40 °C above the first transition, for DsbB under LMNG, DM, and LDAO conditions. The measured thermostability for DsbB in detergent micelles from most stable to least stable is DDM > LMPG > LMNG > DM > LDAO. The fraction unfolded curves (baseline corrected) for each detergent condition are reported in **Figure S4** for visualization of the T_m_ values at the inflection point as well as the dynamic light scattering data collected as a cumulative radius plot in **Figure S3A-E (third row)**.

### The experimentally determined T_m_ value for the fluorescent MSP-based nanodisc is well outside the range of T_m_ values observed for DsbB

We also tested the thermostability of a fluorescent MSP-based nanodisc to ensure that it was not contributing to the three-state/biphasic unfolding pattern observed for DsbB. **Figure 3A** shows the F350/F330 ratio for the unfolding curve for a fluorescent MSP-based nanodisc in the absence of DsbB. **Figure 3B** shows the respective experimental derivative plot for the unfolding curve. The inflection point corresponds to a T_m_ value of 86.6 °C, which is more than 10 °C above the melting temperatures observed for DsbB in a dark nanodisc (see – **Figure 2B**). **Figure S1B** shows the fraction unfolded curve (baseline corrected) for the fluorescent MSP-based nanodisc for visualization of the T_m_ value at the inflection point. This data illustrates that the baseline DSF trace from fluorescent nanodiscs would be problematic had these nanodiscs been used for this study of DsbB. The dynamic light scattering data collected for the empty fluorescent nanodiscs is shown in **Figure S2C (third row)** as a cumulative radius plot indicating that aggregation of the empty fluorescent nanodiscs falls outside the range of T_m_ values observed for DsbB in the dark nanodisc.

### The engineered *dark loop* DsbB construct does not appear to change the three-state /biphasic unfolding pattern for DsbB but displays higher transition temperatures

Otzen *et al*. reported that a single exponential for both the kinetic folding and unfolding of DsbB in mixed micelles was observed suggesting that DsbB refolds/unfolds as a single cooperative unit with a strong coupling between the four transmembrane domains and periplasmic loop regions^20^. We were interested in determining if this cooperativity of unfolding was specific to detergent conditions considering that our thermostability measurements of DsbB in a nanodisc revealed a biphasic unfolding pattern. We hypothesized that under nanodisc conditions, the three-state/biphasic pattern was the result of the extramembrane loop region unfolding independent of the transmembrane core. To test this hypothesis, we engineered a *dark loop* DsbB construct where the Trp residues in the 46-residue extracellular loop region were mutated out to spectroscopically silence this region of the protein (i.e., W113F, W119L, and W135F). **Figure S5A** shows the F350/F330 ratio for the unfolding curve for the *dark loop* construct assembled in a dark nanodisc. **Figure S5B** shows the respective experimental derivative plot for the unfolding curve revealing two inflection points at 85.3 °C and 91.7 °C. Indeed, nanoDSF measurements of *dark loop* DsbB reconstituted in a dark nanodisc showed that the protein still undergoes a three-state/biphasic unfolding process, consistent with the fact that DsbB unfolds as a cooperative unit (see – **Figure S5**). This observation further supports that an early unfolding of the periplasmic loop regions is not a contributing factor to the observed biphasic unfolding pattern. However, we note the melting temperatures observed for the *dark loop* construct are near or above those observed for the empty fluorescent nanodisc (i.e., 86.6 °C). **Figure S1C** shows the fraction unfolded curve (baseline corrected) for the *dark loop* DsbB construct reconstituted in a nanodisc for visualization of the T_m_ value at the inflection point. Additionally, we report the dynamic light scattering data that we collected for the *dark loop* DsbB construct in dark nanodiscs as a cumulative radius plot in **Figure S2E (third row)**.

### The thermal unfolding mechanism for DsbB measured under reducing conditions shifts the unfolding curve from three-state/biphasic to two-state/monophasic

We purified wild-type DsbB under reducing conditions (i.e., the addition of 2 mM TCEP) to test the hypothesis that the interloop disulfide bond between cysteine residues C41 and C130 plays a key role in the formation of the intermediate unfolding state. **Figure 4A** shows the F350/F330 ratio for the unfolding curve for C41-C130 reduced DsbB assembled in a dark nanodisc. **Figure 4B** shows the respective first derivative plot for the unfolding curve revealing a single inflection point corresponding to a T_m_ value of 76.6 °C, which is within the range of the T_m2_ value observed for DsbB under non-reducing conditions. This suggests that the intermediate unfolding state observed for DsbB is mediated by the C41-C130 disulfide bond. More interestingly, this distinct three-state/biphasic unfolding pattern is not observed under detergent conditions further suggesting that this interloop disulfide bond intermediate state is mediated by the presence of lipids. **Figure S1D** shows the fraction unfolded curve (baseline corrected) for C41-C130 reduced DsbB reconstituted in a nanodisc for visualization of the T_m_ value at the inflection point and **Figure S2B (third row)** shows dynamic light scattering data collected for DsbB under reducing conditions in a dark nanodisc as a cumulative radius plot.

**Figure 4.**
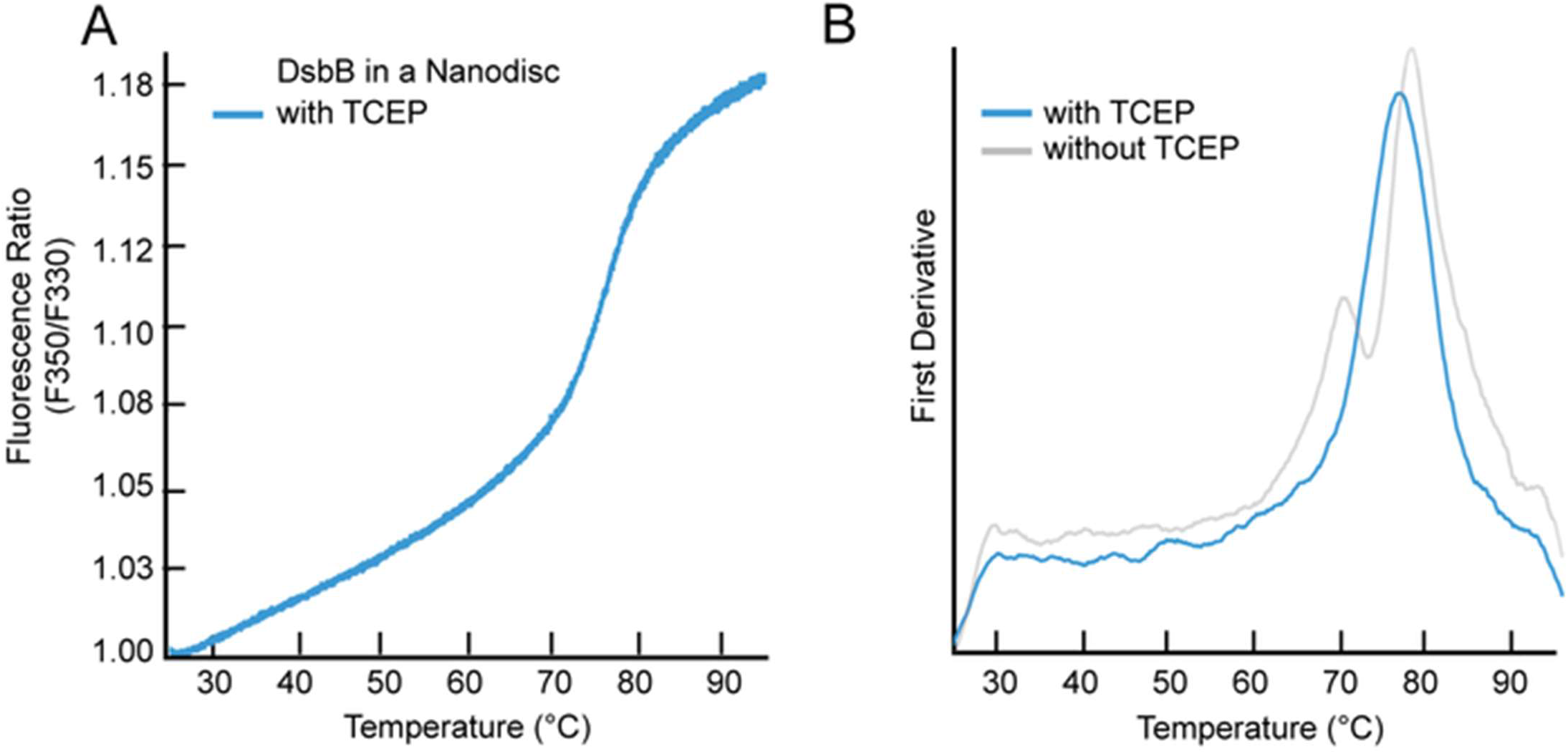
NanoDSF measurements for DsbB reconstituted in dark nanodiscs under reducing conditions. **(A)**. The F350/F330 thermal unfolding curve of DsbB reconstituted in the dark nanodisc in the presence of the reducing agent TCEP at 2 mM. (**B**) The first derivative plot shows a single inflection point corresponding to a T_m_ value of 76.6 ± 0.5 °C. The first derivative plot for DsbB reconstituted in the dark nanodisc in the absence of reducing conditions has been overlayed as a comparison. All samples were run in triplicate and the average is plotted.

## Discussion

### The intermediate minor transition state observed for the thermal unfolding of DsbB in a dark nanodisc is absent under detergent micelle conditions

Tools to evaluate membrane protein stability in the presence of lipids or a lipid-based model membrane system remain a technical challenge and are often reverted to studies in detergent micelles. We describe in this work the development and evaluation of dark nanodiscs as a model membrane system for nanoDSF thermostability measurements. We present this as a new tool that can be used to evaluate how lipid-protein interactions regulate membrane protein stability. We show that the dark nanodiscs are fluorescently undetectable by nanoDSF, making them an ideal model membrane system to investigate the effect of lipids and specific lipid types on membrane protein thermostability.

We observed that DsbB reconstituted in a dark nanodisc follows a three-state/biphasic thermal unfolding model that can be described in terms of a native state (**N**), an intermediate state (**I**), and an unfolded state (**U**). **Figure 5** depicts the assembled DsbB-dark nanodisc complex as it unfolds along the thermal gradient. In this model, the DsbB thermal unfolding curve does not reach a plateau (see – **Figure 2A**) suggesting that residual native-like secondary structural elements are maintained in the unfolded state. The thermostability profile for DsbB in nanodiscs compared to detergent micelles revealed two distinct differences – (**i**) the thermostability for DsbB in a nanodisc is significantly higher than under micelle conditions and (**ii**) the distinct three-state/biphasic unfolding model of DsbB in a nanodisc switches to a two-state/monophasic process under micelle conditions revealing a major thermal transition state. These observations further signify the impact and role lipids play in the thermal stability and thermal unfolding of membrane proteins.

**Figure 5.**
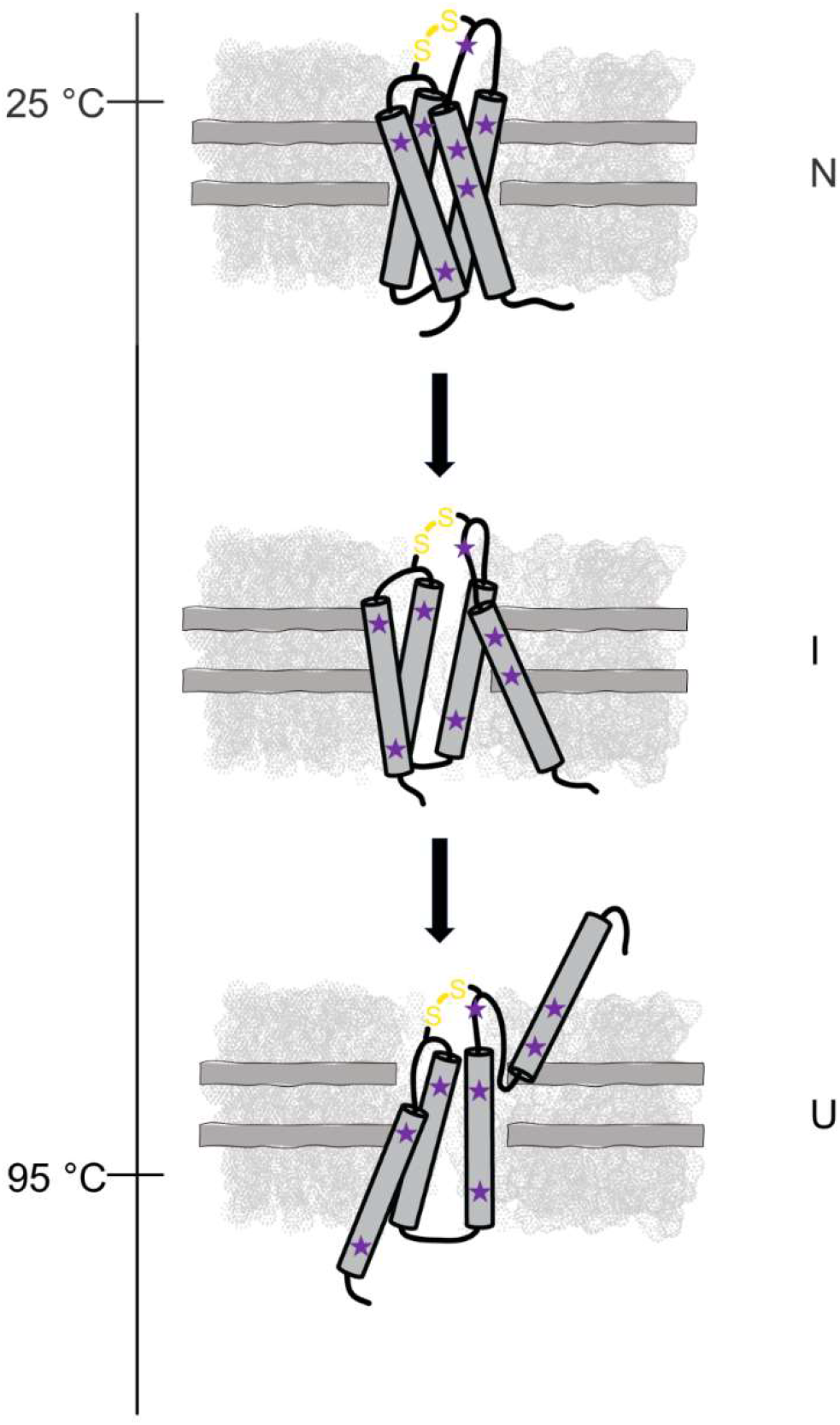
Schematic summarizing the proposed three-state unfolding model for DsbB in a dark nanodisc. DsbB reconstituted in a dark nanodisc system loaded with DMPC and DMPG lipids at a 3:2 molar ratio and the non-fluorescent MSP as it thermally unfolds from the native state (**N**) to an intermediate state (**I**) and to the unfolded state (**U**). We propose that the three-state/biphasic model is the result of a disulfide-bond mediated intermediate state in the presence of lipids. DsbB is shown in dark grey, the dark nanodisc system is shown in light grey, the tryptophan and tyrosine residues are shown as purple spheres, and the C41-C130 interloop disulfide bond is shown in yellow.

The decrease in thermal stability for detergent-solubilized DsbB in micelles compared to DsbB reconstituted in nanodiscs is not surprising. It is well-recognized that the function and stability of membrane proteins are compromised during isolation from the membrane and delipidation through detergent solubilization and even more, can be restored by reconstituting the protein back into a lipid-based model membrane system. For example, the light-harvesting chlorophyll *a/b* complex (LHCIIb) of photosystem (PS) II was reported to have a lower thermostability in DDM micelles (T_m_ = 59.4 °C) and when reconstituted in proteoliposomes, increased to T_m_ values as high as 74.9 °C in the presence of monogalactosyl diacylglycerol (MGDG) lipids^23^. Notably, the thermal stability for LHCllb was determined over a range of lipid compositions and revealed that specific lipid types can shift the T_m_ value as much as 15 °C (i.e., phosphatidylglycerol vs. MGDG)^23^. Thermograms for the Na,K-ATPase cation transporter reconstituted in DPPC (dipalmitoylphosphatidylcholine) and DPPE (dipalmitoylphosphatidylethanolamine) proteoliposomes revealed different thermostability profiles under varying concentrations of cholesterol suggesting that the mechanism by which the protein thermally unfolds is dependent on cholesterol content^24^.

The three-state/biphasic unfolding model of DsbB in a nanodisc is consistent with prior observations of the folding/unfolding of DsbB in mixed micelles measured by stopped-flow kinetic experiments^20,25^. Although thermal denaturation of membrane proteins is known to be an irreversible process, and thus true ΔG values cannot be extrapolated from this data, we utilized the two-state/monophasic fitting model in MoltenProt to calculate an estimated value of 3.8 ± 0.1 kcal/mol for DsbB in DM detergent micelles. We used this as a point of comparison to prior kinetic studies of DsbB where they determine a true ΔG value in DM detergent micelles of 4.2 ± 1.4 kcal/mol. However, in our studies, detergent-solubilized DsbB in DM micelles appears to exhibit a two-state/monophasic thermal unfolding behavior with a major transition in contrast to the three-state/biphasic unfolding reported by Otzen *et al*. There is a second less-defined peak that occurs 40 °C post the first thermal transition and is discussed further below. Furthermore, we observe this same behavior in the case of LMNG and LDAO micelles (see – **Figure S3C and Figure S3E)**. In one published study, the erythropoietin-producing hepatocellular carcinoma B receptor (EphB) kinase domains were investigated by thermal unfolding and chemical denaturation experiments and similarly, the authors also observed a two-state/monophasic and three-state/biphasic unfolding behavior for each respective method^26^. One potential explanation for these unfolding mechanism differences is that thermal denaturation experiments are prone to irreversible aggregation whereas chemical denaturation with urea is reversible and does not involve protein aggregation.

To deconvolute the second observed peak for DsbB under DM, LMNG, and LDAO conditions we evaluated the dynamic light scattering (DLS) traces collected in parallel with the fluorescence measurements (**Figure S3A-E**), which present several interesting observations. First, the cumulative radius plots for DsbB in DM, LMNG, and LDAO each show a significant increase in system size (i.e., >4000 nm) towards the end of the thermal gradient. This corresponds to the second less-defined peaks in the first derivative plots. This suggests that the delay in the onset between the first and second peaks is the result of protein aggregation and micelle dissociation from the unfolded protein state. Second, the cumulative radius plots for DsbB in detergent micelles show marked variability. DM, LMNG, and LDAO have similar readouts with a drastic increase in system size at the end of the thermal gradient. In contrast, DDM and LMPG both show a minimal increase in the cumulative radius at the end of the thermal ramp. We speculate the different patterns observed for the change in system size are attributed to the different chemical and physical properties of the detergent micelles. For example, DsbB has a +4 charge state at the pH of our measurements, and detergents such as LMPG are negatively charged^27^.

We propose that electrostatic interactions between LMNG and DsbB are strong enough to prevent micelle dissociation from the unfolded state and thus, prevent aggregation.

In the case of DsbB in a dark nanodisc, the cumulative radius plot in **Figure S2A (third row)** shows that the system size increases moderately to 340 nm at a temperature of 76.5 °C and 664 nm at a temperature of 83.9 °C. These temperature-dependent changes in cumulative radius occur after the minor and major thermal unfolding states observed for DsbB. This information suggests that (i) the change in system size correlates with the unfolding of DsbB and/or (ii) protein unfolding at the midpoint melting temperatures is followed by protein aggregation.

### The periplasmic loops and transmembrane domain regions of DsbB reconstituted in a nanodisc unfold as a cooperative unit along the thermal gradient

The experimental derivative plot for the unfolding curve of *dark loop* DsbB in a dark nanodisc revealed two inflection points at 85.3 °C and 91.7 °C, which are in close range and above the melting temperature observed for the empty fluorescent nanodisc (i.e., 86.6 °C). Despite the apparent increase in melting temperature, the *dark loop* construct still exhibits a three-state/biphasic unfolding pattern suggesting that DsbB unfolds as a cooperative unit. To determine if the apparent increase in stability of the *dark loop* DsbB construct compared to the wt construct was due to a stabilizing effect of the *dark loop* mutations, we performed Rosetta calculations on a DsbB NMR ensemble to estimate the relative energy between the two constructs. We found no statistically significant difference between wt and *dark loop* DsbB energy in terms of Rosetta Energy Units (REU) (**Figure S6**). This suggests that the *dark loop* mutations do not stabilize the structure enthalpically. Although outside the scope of this work, another potential explanation for this increase in thermal stability for the *dark loop* construct could be that we expressed it in nutrient-rich Luria-Bertani (LB) media as opposed to minimal (M9) media in order to obtain enough protein yield for nanodisc reconstitutions. LB media is known to be higher in ubiquinone content due to the presence of additional precursors required for synthesis^28^ and ubiquinone is known to bind and structurally stabilize DsbB^29^. It is also possible that interactions between the engineered Phe residues and lipids in the nanodisc caused a shift in the melting temperatures and enhanced stability.

### Lipid-mediated interactions between the catalytically conserved cysteine residues of DsbB and the dark nanodisc drive the biphasic unfolding mechanism

Time-resolved kinetic experiments showed the stability of DsbB in mixed micelles was reduced by half upon the reduction of the two periplasmic disulfide bonds between residues C41-C44 and C104-C130^20^. Disulfide bonds have been shown to play key roles in membrane protein stability for a number of systems such as G protein-coupled receptors (GPCRs)^30^ and ATP-binding cassette (ABC) transporters^31^. For example, a simulated thermal unfolding of the GPCR rhodopsin revealed that the C110–C187 disulfide bond is key to retinal binding and stabilizing the folded state of rhodopsin^30^. In another example, the intramolecular disulfide bond C592–C608 located in the extracellular loop of the ABC transporter ABCG2 was shown to be critical for protein stability (i.e., plasma localization and expression levels), which was significantly reduced when this disulfide bond was mutated out with glycine residues^31^.

Prior studies have investigated disulfide bond interactions in the presence of lipids that contribute to the stability of proteins. For example, a Raman spectroscopy study of a lung surfactant peptide B co-solubilized in a lipid mixture of dipalmitoyl phosphatidylcholine (DPPC) and PG1,2-dioleoyl-sn-glycero-3-phospho-(1’-rac-glycerol) (DOPG) at a 4:1 lipid to protein ratio suggests that the presence of lipids “locks” the disulfide bond into a distinct conformation^32^. Furthermore, molecular dynamics (MD) simulations have investigated the role of particular disulfide bonds in the mechanism of DsbB as a disulfide bond formation mediator^33^. These MD simulations reveal that DsbB undergoes several disulfide bond rearrangements resulting in conformational changes in the periplasmic loop region. Specifically, the loop adopts a compact conformation to accommodate the C41-C130 disulfide bond. This suggests that under reducing conditions, DsbB cannot adopt this compact conformation in the periplasmic loop region, leading to a lack of stabilization and loss of the intermediate state. Furthermore, the intermediate state is only observed under lipid nanodisc conditions, suggesting that lipid-mediated interactions between the cysteine residues drive the three-state/biphasic thermal unfolding pattern.

In this work, the construct used has a disulfide bond between residues C41 and C130 and is referred to as the interloop disulfide-bond intermediate state. We find that under reducing conditions (2 mM TCEP), the thermal unfolding pattern for DsbB shifts to a two-state model and a loss of the first minor thermal transition as shown in **Figure 4B** (i.e., T_m1_ = 70.5 °C). We show that the second major thermal transition state (T_m2_) is maintained under reduced conditions. From these results, we conclude that (i) the interloop disulfide bond plays a critical role in the thermal unfolding of DsbB and (ii) the presence of lipids contributes to the observed minor thermal transition state as this is not observed under detergent conditions. In summary, this suggests that the disulfide bond formed between the catalytically conserved cysteine residues contributes to the stabilization of a metastable intermediate state during the thermal unfolding of DsbB.

### Outlook

Overall, the method developed herein will be broadly applicable in membrane protein biophysics research. It provides a simple tool to dissect the effect of lipid-protein interactions on membrane protein thermostability – interactions that are typically missed or difficult to measure using existing methods. For example, genetic mutations lead to the loss of important intermolecular interactions that cause structural instability and misfolding of proteins. This leads to serious genetic diseases such as cystic fibrosis (CF), long QT syndrome, and Charcot-Marie tooth disease. Specifically, these diseases are caused by the misfolding of an integral membrane protein including the CF transmembrane conductance regulator (CFTR)^34^, the KCNQ1 ion channel^35^, and the peripheral myelin protein 22 (PMP22)^36^, respectively. In such cases, mutations compromise membrane protein stability, resulting in aberrant protein function. Experimental and high-throughput measurements are key for understanding (i) the effects of genetic mutations on the thermostability of membrane proteins and (ii) how mutations reshape native lipid-protein interactions to drive disease states, which have been historically underappreciated^37,38^. Furthermore, our method will be used to test membrane protein design candidates in cases where the goal is to design a more thermostable protein to facilitate biochemical or structural studies.

## Acknowledgments

This work was supported by NIH grants R01 GM080403, R01 HL122010, and R01 GM129261. The authors further acknowledge funding by the Deutsche Forschungsgemeinschaft (DFG) through SFB1423, project number 421152132. Jens Meiler is supported by a Humboldt Professorship of the Alexander von Humboldt Foundation. EFM was supported by a predoctoral fellowship from the National Heart, Lung, and Blood Institute (F31 HL162483-01A1). We also thank Dr. Vadim Kotov for technical assistance and discussions regarding the use of MoltenProt for data visualization and interpretation. This work was conducted using the resources of the Center for Structural Biology at Vanderbilt University. We especially thank the Lab of Dr. Charles Sanders for technical assistance with the nanoDSF experiments and insightful discussions. The nanoDSF instrument used in this work was acquired via an ANCORA grant to the Sanders lab from Deerfield Drug Discovery (3DC). This work was also conducted using the resources of the Advanced Computing Center for Research and Education (ACCRE) at Vanderbilt University.

## Author Contributions

Conceptualization, KVL. Data curation, JAS and JRM; Formal analysis, JAS, JRM, KVL; Funding acquisition, JM; Investigation, JAS, JRM, EFM, KVL; Methodology, KVL; Supervision, KVL and JM.; Writing—original draft, JAS and EFM; Writing—review and editing, EFM, JM, and KVL. All authors have read and agreed to the published version of the manuscript.

## Competing Interests

The authors declare no competing interests.

## Methods

### Expression and purification of the non-fluorescent membrane scaffold protein (MSP1D1) and fluorescent MSP1D1 construct

The non-fluorescent and fluorescent MSP1D1 protein genes were cloned in a pET28a vector containing a tobacco etch virus (TEV) protease-cleavable N-terminal His_6_ tag. BL21(DE3) competent *E. coli* cells were transformed with the target plasmid and grown on Luria-Bertani (LB) agar plates overnight at 37 °C. A single colony was selected and used to inoculate a 150 mL LB starter culture and grown overnight at 37 °C. Flasks containing 1L of terrific broth (TB) media were inoculated with 10 mL of the starter culture and grown at 37 °C at 230 rpm in the presence of 100 ng/ml kanamycin. Protein expression was induced at an OD_600_ of 0.6-0.8 with 1 mM IPTG (isopropyl β-D-1-thiogalactopyranoside) and cultured for another 24 hours at 20 ºC. Cells were harvested by centrifugation at 6500 rpm for 20 minutes and cell pellets were stored at -80 ºC or immediately resuspended in lysis buffer for purification.

Cell pellets expressing the non-fluorescent or fluorescent MSP1D1 proteins were resuspended (5 mL/gram of pellet) in Buffer A (50 mM Tris, 300 mM NaCl, pH 8.0) with the addition of 1 mM PMSF, 1% Triton X-100, EDTA-free protease cocktail inhibitor tablet (Sigma-Aldrich) (1 tablet/per 50 mL of buffer), and 5 mM Mg acetate. The cells were lysed by sonication for 10 minutes (60% amplitude with 5 seconds on/5 seconds off) on ice. The cell lysis solution was centrifuged at 50,000 x g for 20 minutes and the supernatant was loaded onto a Ni-NTA gravity-flow column. The Ni-NTA resin was washed with 10 column volumes of the following in this order: Buffer A and 1% Triton X-100, Buffer A and 75 mM sodium cholate, Buffer A, and Buffer A plus 20 mM imidazole. Proteins were eluted with Buffer A containing 500 mM imidazole. Isolated proteins were dialyzed against TEV cleavage buffer (20 mM Tris-HCl, 100 mM NaCl, 1 mM DTT, pH 7.5) at 4 ºC for 24 hours. The TEV cleavage reaction was set up using TEV protease at a 1:50 ratio of protease to protein to remove the N-terminal His_6_ tag.

The cleaved sample was loaded onto a second Ni-NTA column to remove the TEV protease and N-terminal His_6_ tag. The flow-through was collected containing the target protein. The MSP1D1 samples were size excluded using a 120 mL HiLoad® 16/600 Superdex® S200 preparative column (Cytiva) equilibrated with reconstitution buffer (40 mM Tris-HCl, 200 mM NaCl, pH 7.5). Protein fractions were analyzed by SDS-PAGE. Pooled fractions were quantified using a Bradford assay and concentrated to 500 μM in an Amicon centrifugal filter (10,000 MWCO) and analyzed by SDS-PAGE to confirm the purity of the samples were greater than 95%. As a control, the Ni-NTA bound TEV protease and His_6_ tag were eluted from the resin. The resin was washed with reconstitution buffer followed by a 10 mM imidazole wash. The TEV protease and cleaved His_6_ tag were eluted with reconstitution buffer and 500 mM imidazole.

### Expression and purification of DsbB

The “wild-type” (wt) DsbB protein construct^29^ was inserted into pET22b vector and transformed into BL21(DE3) *E. coli* cells. Cells were grown on Luria-Bertani (LB) agar plates containing 100 μg/mL ampicillin overnight at 37 ºC. A single colony was selected and used to inoculate a 150 mL starter culture grown overnight at 37 ºC. Flasks containing 1L of minimal (M9) media were inoculated with 10 mL of starter culture and grown in the presence of 100 μg/mL ampicillin. Cells were cultured at 37 °C at 230 rpm until the OD_600_ reached 0.6-0.8. Cells were transferred to room temperature and protein expression was induced with 1 mM IPTG for 18 hours. Cells were harvested by centrifugation at 6500 rpm for 15 minutes at 4 ºC. Cell pellets were stored at -80 ºC or immediately resuspended in lysis buffer for purification.

Each cell pellet was resuspended in 40 mL of Buffer B (50 mM Tris-HCl, 300 mM NaCl, pH 8.0) with the addition of 1 mM PMSF, 5 mM Mg Acetate, and 1 EDTA-free protease cocktail inhibitor tablet (Sigma-Aldrich) (1 tablet/per 50 mL of buffer). The cell lysis solution was rotated at 4 °C for 30 minutes and then lysed by sonication for 10 minutes (60% amplitude with 5 seconds on/5 seconds off). The membrane fraction was pelleted by ultracentrifugation at 100,000 x g for 1 hour at 4 °C and the supernatant was discarded. The membrane fraction was homogenized on ice with Buffer B (20 mL/pellet) containing 1% dodecylphosphocholine (DPC, FOS-CHOLINE-12, Anatrace). Homogenized cells were ultracentrifuged at 100,000 x g for 30 minutes to remove the insoluble fraction. The remaining supernatant was loaded onto a Ni-NTA gravity-flow column equilibrated with Buffer B and 0.05% DPC. The Ni-NTA column was washed with Buffer B and 0.05% DPC followed by Buffer B plus 0.05% DPC and 10 mM imidazole. DsbB was eluted with Buffer B plus 0.05% DPC and 500 mM imidazole. The collected sample was concentrated to 500 μM and size excluded using a 120 mL HiLoad® 16/600 Superdex® S200 preparative column (Cytiva) equilibrated with FPLC buffer (50 mM Tris-HCl, 300 mM NaCl, 0.15% DPC, pH 8.0). The pooled fractions were concentrated to 500 μM in an Amicon centrifugal filter (10,000 MWCO) and analyzed by SDS-PAGE to confirm the purity of the sample was greater than 95%. For the reduced DsbB sample, we followed the above protocol with the exception that each buffer in the purification protocol was supplemented with 2 mM TCEP.

### *Dark loop* DsbB mutagenesis, expression, and purification

A modified DsbB protein construct containing mutations W113F, W119F, and W135F was engineered and inserted into a pET22b vector. The plasmid was transformed into BL21(DE3) *E. coli* cells and expressed and purified following the same protocol as described for wt DsbB. The protein yield for this construct was low. To increase the protein yield of this *dark loop* construct we performed a multiple sequence alignment (MSA) and identified that residue W119 is highly conserved. For this reason, we engineered a second construct with a W119L mutation in place of the W119F mutation using site-directed mutagenesis. This MSA-modified construct containing mutations W113F, W119L, and W135F was transformed into C43(DE3) *E. coli* cells and expressed in LB media to optimize the expression of the *dark loop* construct. Flasks containing 1L of LB media were inoculated with 10 mL of starter culture and grown in the presence of 100 μg/mL ampicillin. Cells were cultured at 37 °C at 230 rpm until the OD_600_ reached 0.6-0.8 and protein expression was induced with 1 mM IPTG for 5 hours at 37 °C. The purification of *dark loop* DsbB followed the same protocol as described above for wt DsbB.

### Reconstitution reactions for DsbB in dark nanodiscs and reconstitution reactions for empty nanodiscs

All purified proteins were concentrated to 500 μM as described above and reconstitutions were set up as previously described by Nasr. *et al*^39^. Lipids were prepared as an 80 mM stock of DMPC and DMPG lipids at a 3:2 molar ratio solubilized in 10% DM. Bio-beads SM2 resin (Bio-Rad) was prepared freshly prior to each reaction by washing the beads with 10 mL of methanol followed by excess deionized water. DsbB was assembled into dark nanodiscs by adding DsbB (160 μM), non-fluorescent MSP1D1 (160 μM), and lipids (13.6 mM) to reconstitution buffer at a 1:1:85 ratio, respectively. Reaction components were added to the reconstitution buffer in the order of lipids, MSP1D1, and DsbB. The reaction was then equilibrated at room temperature for 1 hour. Fresh Bio-beads (0.5g of wet beads/300 μl of total reaction volume) were added to the reaction and allowed to tumble overnight at room temperature to remove all detergents. The DsbB-containing nanodisc solution was collected to remove the Bio-beads and loaded three times onto a Ni-NTA column equilibrated with reconstitution buffer (i.e., DsbB is His_6_-tagged). The Ni-NTA resin was washed with excess reconstitution buffer to remove empty nanodiscs and/or large vesicles. The assembled DsbB-nanodisc complex was then size excluded at room temperature using a 24 mL Superdex® 200 Increase 10/300 GL column (Cytiva) equilibrated with reconstitution buffer. Selected fractions were analyzed by SDS-PAGE to confirm the purity of the DsbB-nanodisc samples. Fractions were pooled and concentrated at room temperature using a low spin speed (i.e., 2500 x g) in an Amicon centrifugal filter (30,000 MWCO) to 1 mg/ml. Empty nanodisc reconstitution reactions were performed using the same protocol described above with the exception that the volume of DsbB was replaced with reconstitution buffer (i.e., the reaction volumes were the same for each reconstitution). Reduced DsbB was assembled into nanodiscs as described above with the exception that the reconstitution buffer was supplemented with 2 mM TCEP.

### Preparing detergent-solubilized DsbB under different micelle conditions for nanoDSF measurements

To prepare DsbB under different detergent micelle conditions, we followed the method outlined for a membrane protein stability detergent screen as described^40^. Briefly, DPC-solubilized DsbB was diluted 10-fold into detergent buffers (DDM, DM, LDAO, LMPG, and LMNG) that were made with a 50-fold excess of the critical micelle concentration (CMC) of the target detergent. The diluted DsbB samples were incubated at 4 °C overnight to ensure detergents were fully exchanged. All detergent micelle samples were prepared to a final concentration of 1 mg/mL.

### NanoDSF thermostability measurements

All samples were run on a Prometheus NT.48 nanoDSF instrument (NanoTemper Technologies). Samples were set up in triplicate at a final concentration of 1 mg/mL (10 μL/capillary). The *dark loop* DsbB nanodisc sample was run in triplicate at a final concentration of 2 mg/mL to optimize the signal readout. All nanodisc samples and detergent samples were measured at a scan rate of 0.1 °C per minute over a temperature range of 25 °C to 95 °C and 20 °C to 95 °C, respectively. Samples were excited at 280 nm and the intrinsic tryptophan fluorescence at 350 nm and 330 nm were recorded as a function of temperature to monitor changes upon thermal unfolding.

### Data analysis

All raw nanoDSF data was collected and then exported from the Prometheus Panta Control software for visualization in MoltenProt^40^. The experimental derivatives were exported from MoltenProt and further analyzed in a Jupyter Notebook through Matplotlib^41^. The F350/F330 ratios for each of the triplicate samples were averaged and normalized to the initial value with Matplotlib. The first derivatives were plotted with Matplotlib to determine the inflection points corresponding to the melting temperatures. The fraction unfolded curves were calculated and plotted by fitting the F350/F330 ratio data to a two-state equilibrium model. The rate constants of the unfolding transition and baseline transition as well as the baseline offset and noise were extrapolated. These parameters were utilized to calculate the baseline-corrected experimental curves from the equation utilized by MoltenProt^42^.

### Rosetta calculations

DsbB (PDB ID: 2K73^29^) was broken into its 20 constituent NMR models as described previously^43^. Next, each individual model was used to create a corresponding *dark loop* model by introducing W113F, W119L, and W135F using the MutateResidue mover in Rosetta. Finally, the 20 wt and 20 *dark loop* DsbB models were energetically minimized using FastRelax in Rosetta for one iteration and scored using a membrane scoring function^44^.

## Supplemental Materials

**Table S1.** Table summarizing the measured inflection points (T_m_).

| Experimental Condition | T<sub>m1</sub> (°C) | T<sub>m2</sub> (°C) |
| --- | --- | --- |
| DsbB – dark nanodisc | 70.5 ± 0.3<br>70.0 ± 0.1<br>70.3 ± 0.6 | 77.5 ± 0.1<br>79.1 ± 0.2<br>79.7 ± 0.5 |
| empty fluorescent nanodisc | 86.6 ± 0.6 |  |
| *dark loop* DsbB – dark nanodisc | 85.3 ± 0.6 | 91.7 ± 0.3 |
| DsbB – dark nanodisc + TCEP | 76.6 ± 0.5 |  |
| DsbB – DDM | 55.6 ± 0.2 |  |
| DsbB – LMPG | 45.4 ± 0.1 |  |
| DsbB – LMNG | 41.0 ± 1.2 |  |
| DsbB – DM | 40.1 ± 0.9 |  |
| DsbB – LDAO | 36.7 ± 0.2 |  |

**Figure S1.**
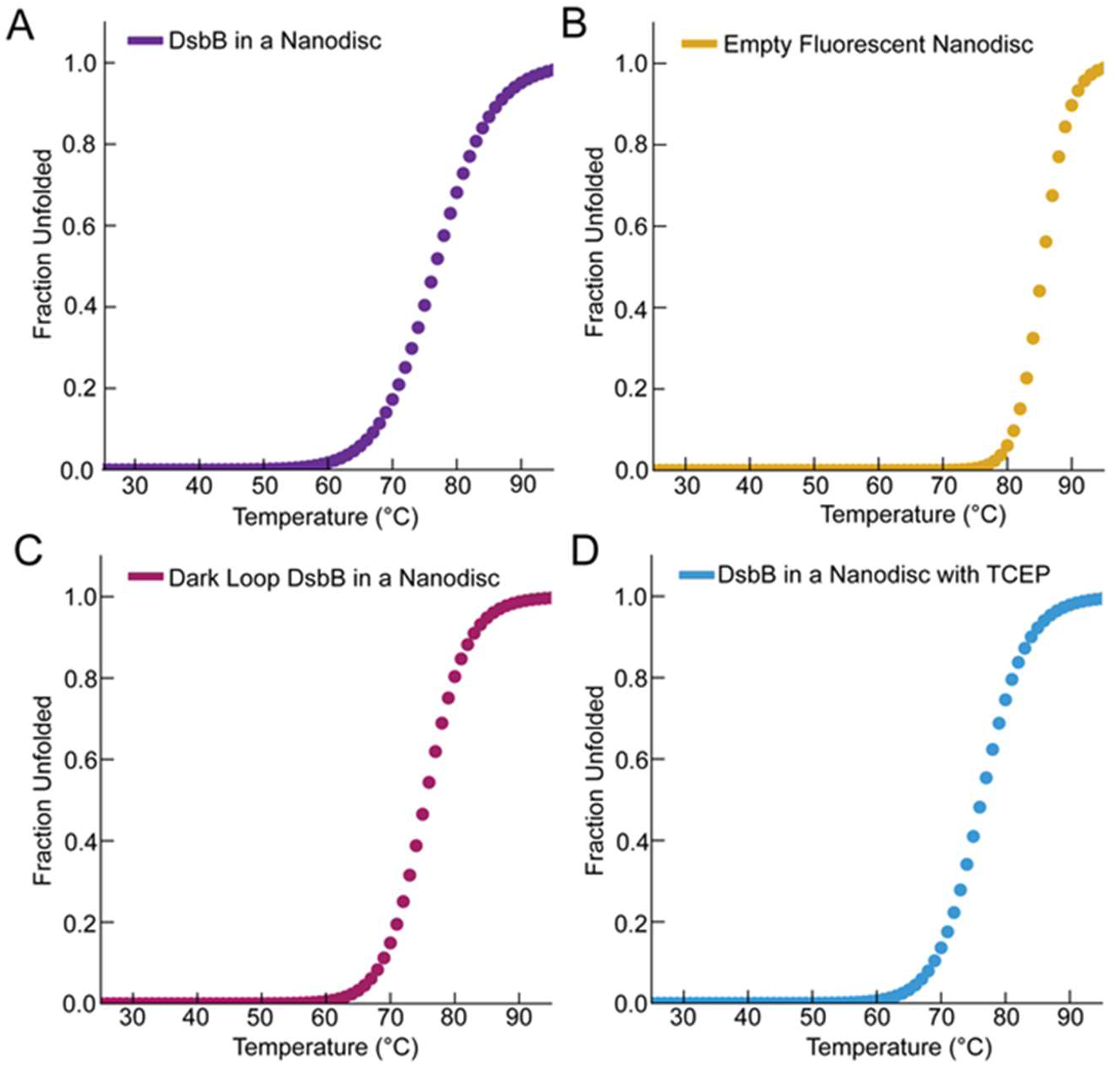
Fraction unfolded plotted as a function of temperature for the nanodisc samples evaluated in this study. For visualization, baseline-corrected experimental curves were calculated from the rate constant of the unfolding transition, the rate constant of the baseline transition, baseline noise, and baseline offset. Fraction unfolded plots are shown for (**A**) DsbB in a dark nanodisc, (**B**) empty fluorescent nanodisc, (**C**) *dark loop* DsbB in a dark nanodisc, and (**D**) DsbB in a dark nanodisc under reducing conditions.

**Figure S2.**
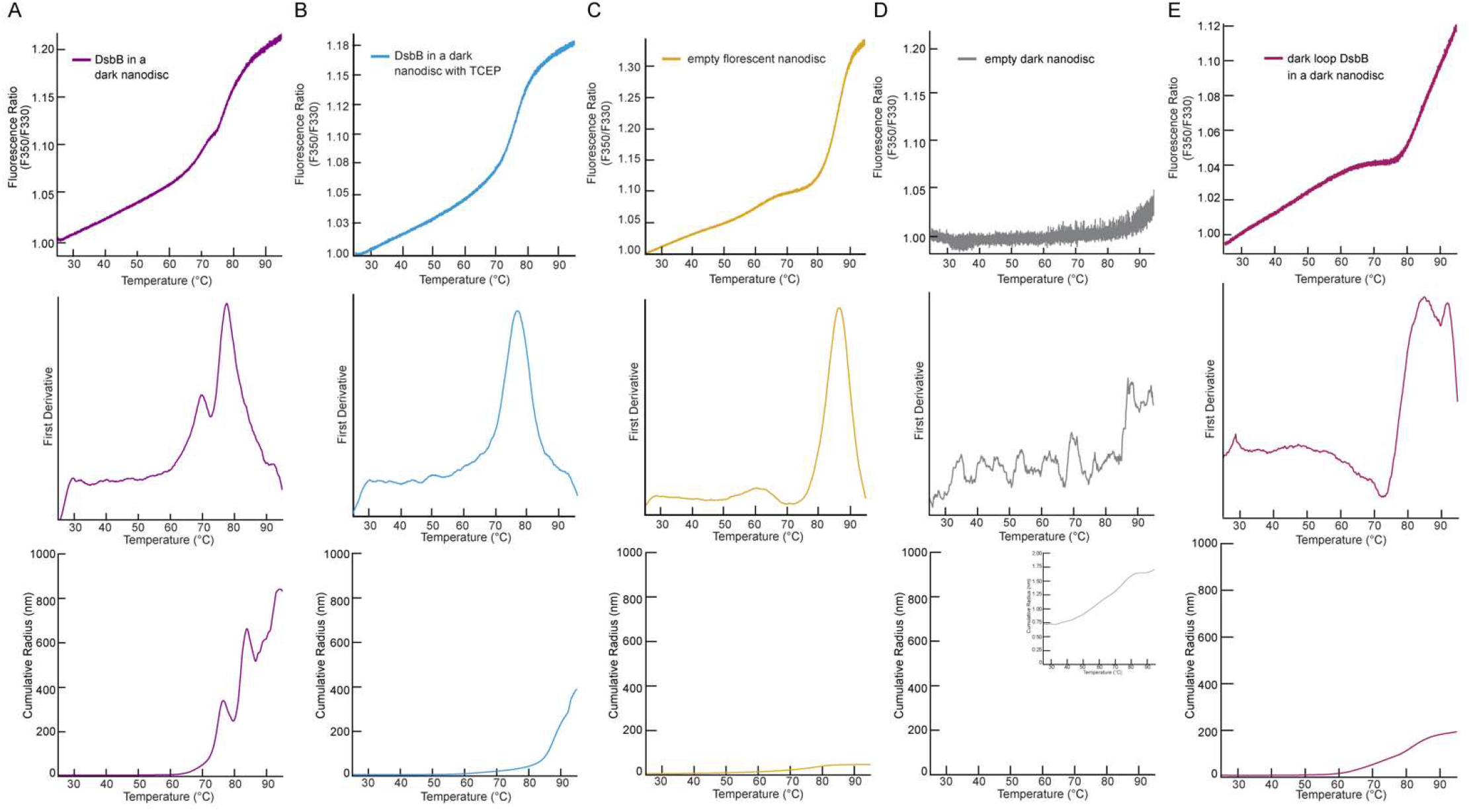
NanoDSF unfolding curves, first derivative, and cumulative radius plots for all nanodisc samples. (**A**) DsbB in a dark nanodisc, (**B**) DsbB in a dark nanodisc under reducing conditions, (**C**) empty fluorescent nanodisc, (**D**) empty dark nanodisc, and (**E**) *dark loop* DsbB in a dark nanodisc. **Top row**: The F350/F330 thermal unfolding curves for each of the nanodisc samples. **Second row**: The first derivative plots for each nanodisc sample. **Third row**: The cumulative radius plots from dynamic light scattering measurements collected in tandem with the fluorescence measurements. All samples were run in triplicate (n=3) with the exception that DsbB in a dark nanodisc was run for three independent sample preparations in triplicate (n=9), the average was plotted.

**Figure S3.**
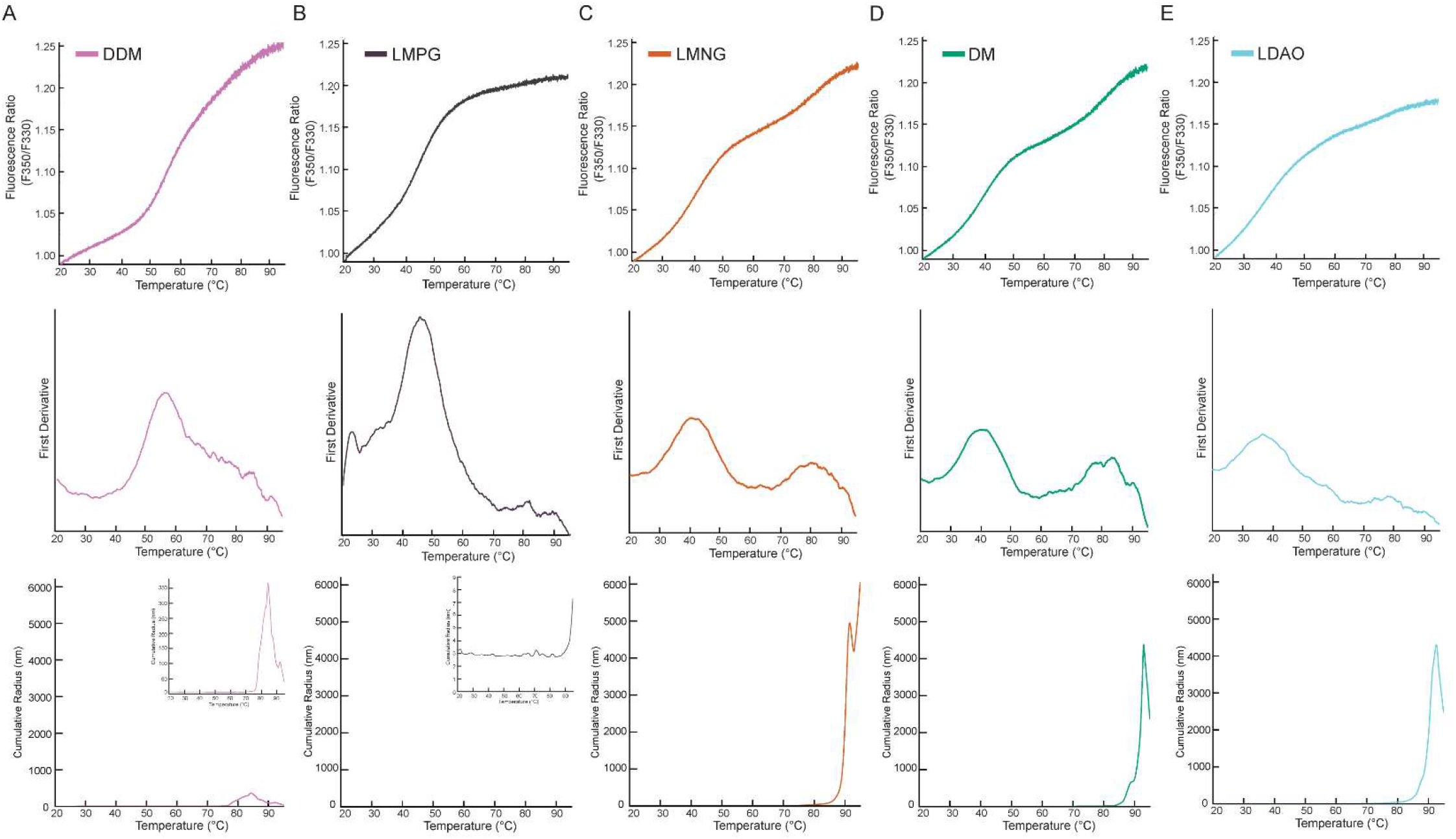
NanoDSF unfolding curves, first derivative, and cumulative radius plots for detergent-solubilized DsbB under a panel of micelle conditions. (**A**) DDM, (**B**) LMPG, (**C**) LMNG, (**D**) DM, and (**E**) LDAO. **Top row**: The F350/F330 thermal unfolding curves for detergent-solubilized DsbB under a panel of different micelle conditions. **Second row**: The first derivative plots for detergent-solubilized DsbB. The inflection points correspond to T_m_ values of 36.7 ± 0.2 °C (LDAO, cyan), 40.1 ± 0.9 °C (DM, green), 41.0 ± 1.2 °C (LMNG, orange), 45.4 ± 0.1 °C (LMPG, purple), and 55.6 ± 0.2 °C (DDM, pink). **Third row**: The cumulative radius plots from dynamic light scattering measurements collected in tandem with fluorescence measurements. All samples were run in triplicate (n=3) and the average was plotted.

**Figure S4.**
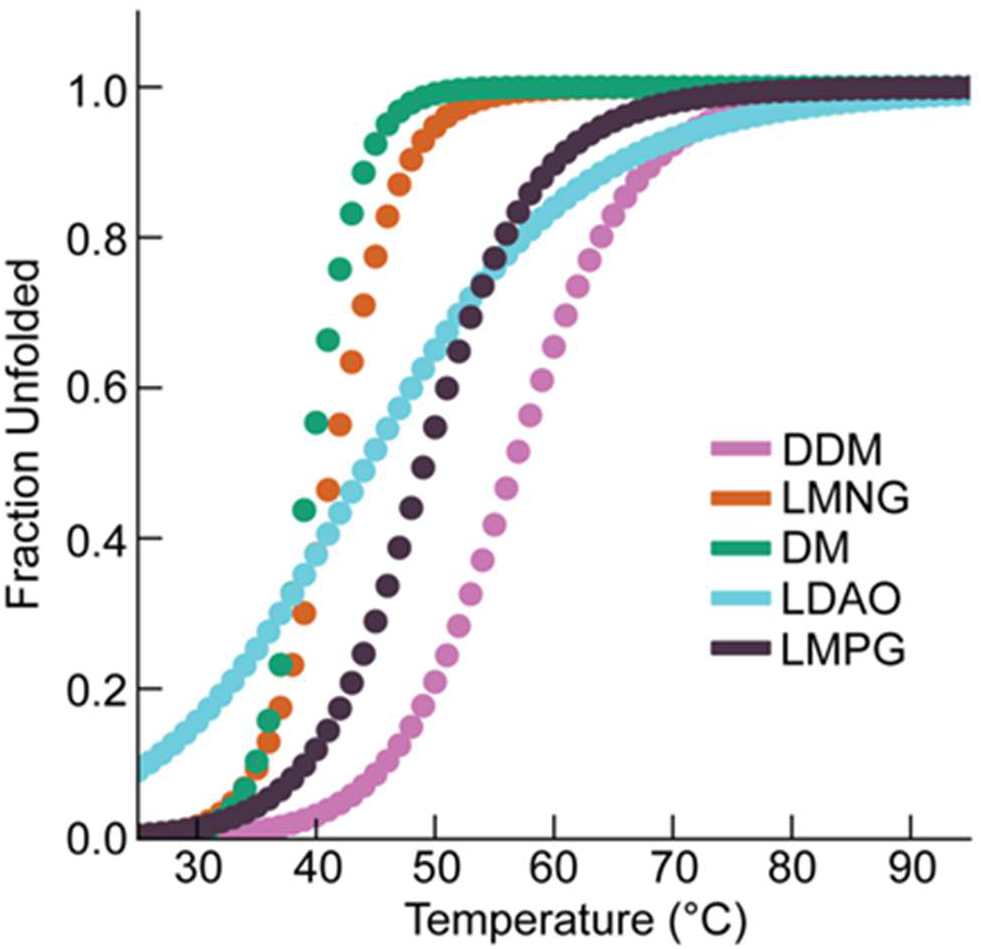
Fraction unfolded plotted as a function of temperature for detergent-solubilized DsbB under a panel of different detergent micelle conditions. The fraction unfolded is plotted for DsbB under a panel of different detergent micelle conditions including DDM, LMNG, DM, LDAO, and LMPG. For visualization, baseline-corrected experimental curves were calculated from parameters derived after fitting the data to an equilibrium two-state unfolding model in MoltenProt^42^.

**Figure S5.**
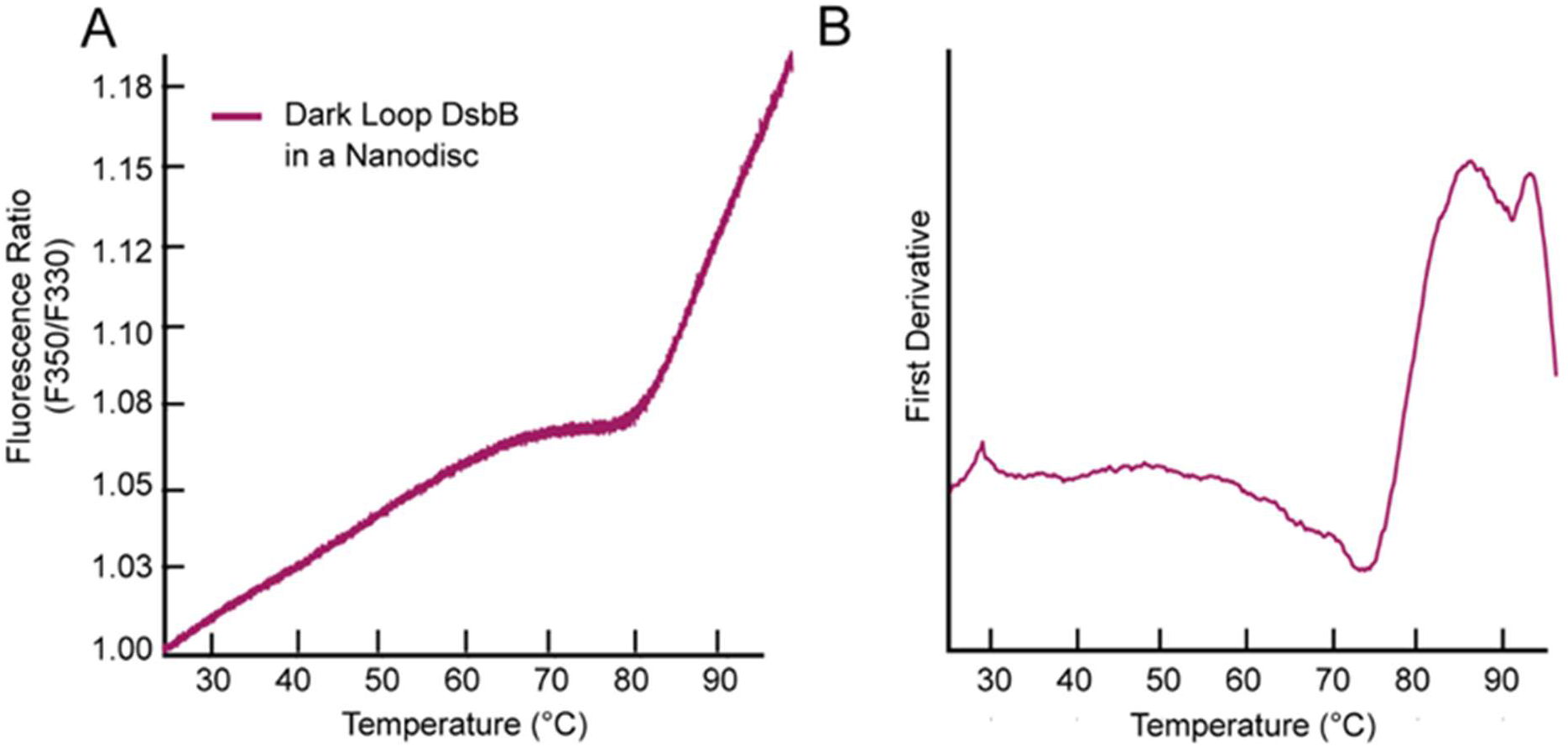
NanoDSF thermal unfolding curve for dark loop DsbB in a dark MSP-based nanodisc. (**A**) The F350/F330 thermal unfolding cure for dark loop DsbB reconstituted in a dark nanodisc model membrane system. (**B**) The first derivative plot of the F350/F330 ratio with respect to temperature for dark loop DsbB in a nanodisc. The inflection points correspond to Tm values of 85.3 °C and 91.7 °C, respectively. All samples were run in triplicate.

**Figure S6.**
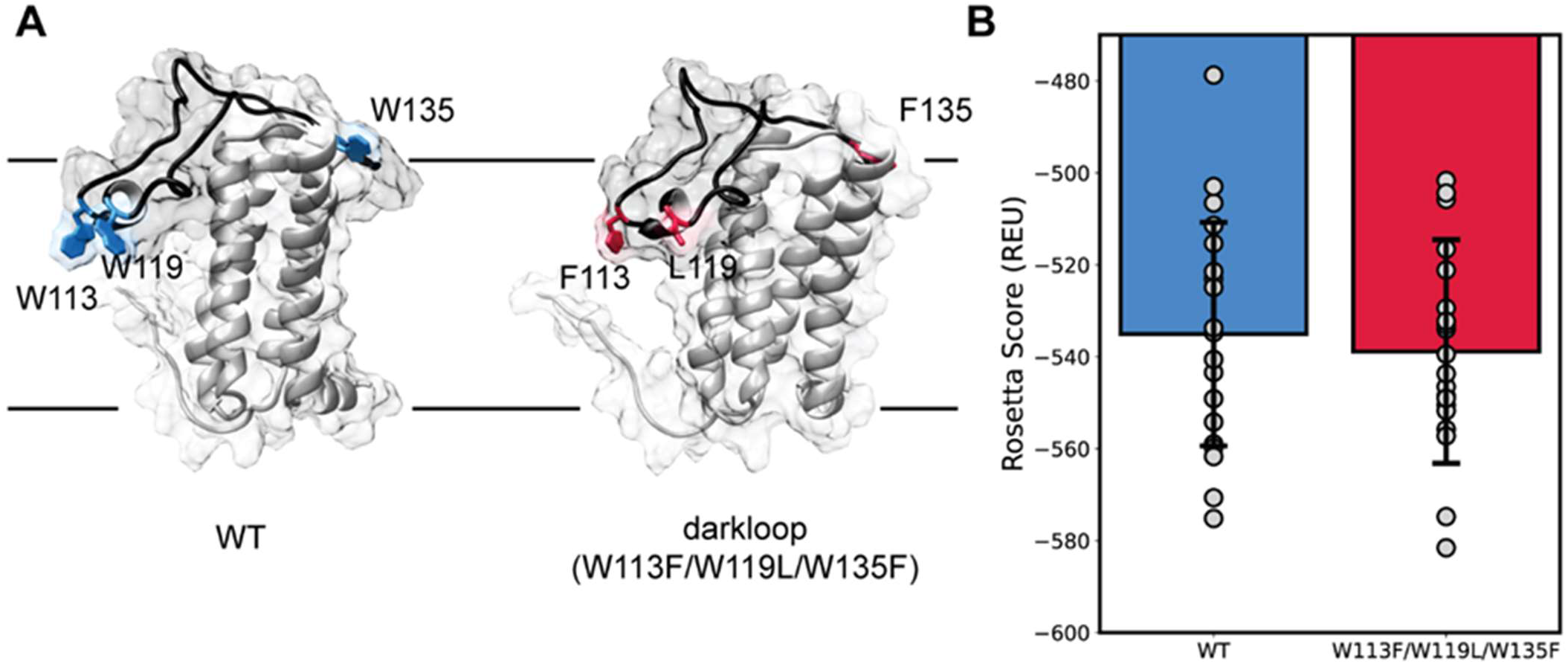
Rosetta calculations of wt and dark loop DsbB based on the ensemble NMR model PDB ID: 2K73. (**A**) Illustration of the residues that were modified to generate the dark loop DsbB construct, which was then relaxed and scored with a Rosetta membrane protein score function. Wt tryptophan residues are depicted in blue and corresponding dark loop residues are depicted in red. The periplasmic loop is shown in black. (**B**) Bar and whisker plot of the membrane score functions for both wt (blue) and dark loop (red) DsbB with a 4 Rosetta energy unit (REU) reduction from -535 REU in wt to -539 REU for dark loop DsbB. Grey dots represent an individual model from the 20-model ensemble and error bars indicate the standard deviations of the scores.

## Notes

### Competing Interest Statement

The authors have declared no competing interest.

